# AMPK Repositions Early Endosomes via Gapex-5 to Promote Delivery of Iron to Mitochondria

**DOI:** 10.1101/2025.10.26.684630

**Authors:** Ayshin Mehrabi, Laura A. Orofiamma, Alyona Ivanova, Kale Allison, Natalie Uzynski, Maria Narciso, Roberto J. Botelho, Eden Fussner-Dupas, Costin N. Antonescu

## Abstract

The regulation of the spatial organization of organelles within cells is critical for coordinating signaling, membrane traffic, and metabolite exchange. Metabolic cues regulate the position and function of lysosomes, yet whether and how metabolic signals may similarly regulate other organelles such as early endosomes (EEs) remains unclear. We find that AMP-activated protein kinase (AMPK), a key regulator of metabolic homeostasis activated in response to nutrient scarcity, triggers movement of EEs to the perinuclear region of cells, leading to enhanced proximity of endosomes to mitochondria and increased delivery of iron to mitochondria. The movement of EEs and increased mitochondrial iron content elicited by AMPK activation requires Gapex-5, a GEF for the early endosome Rab5 previously shown to be an AMPK substrate. These findings reveal a mechanism by which AMPK reprograms endosome positioning to facilitate inter-organelle communication and iron delivery to mitochondria to support metabolic adaptation under conditions of nutrient scarcity.

## Introduction

Eukaryotic cells maintain a highly organized internal architecture, and the spatial positioning of organelles is increasingly recognized as a key determinant of their function. Rather than being randomly distributed, organelles adopt defined subcellular localizations to support efficient signaling, trafficking, and metabolite exchange (Scorrano et al., 2019; Valm et al., 2017). For example, lysosomes exhibit tightly regulated positioning along the microtubule network, with their perinuclear or peripheral distribution influencing mTORC1 activity, autophagic flux, degradative capacity and inflammasome activation (Korolchuk et al., 2011; Pu et al., 2016; Filipek et al., 2017; Matchett et al., 2025). This spatial control is mediated by motor proteins and adaptor complexes that tune lysosome function to nutrient status and cellular stress.

The regulation and functional consequences of early endosome (EE) positioning are less well understood. EEs act as dynamic sorting stations, directing internalized cargo such as transferrin (Tfn) and other receptors toward recycling or degradative fates (Naslavsky and Caplan, 2018; Langemeyer et al., 2018; Yuan and Song, 2020). Early endosomes are classically demarked by the GTPase Rab5, which recruits effectors such as the lipid kinase Vps34 and early endosome antigen 1 (EEA1) to mediate fusion and cargo sorting (Wang et al., 2019; Shin et al., 2005; Gautreau et al., 2014). The position of EEs is controlled by microtubule motors such as dynein-dependent EE movement regulation by FHF complexes that are comprised of Fused Toes (FTS), one of the Hook proteins (Hook1-3), and FTS and Hook-interacting protein FHIP (Abid Ali et al., 2025; Guo et al., 2016; Xiang et al., 2015). The signals that control the position of EEs along microtubule tracks in cells and the consequences of this positioning on endosome function are poorly understood.

One of the better appreciated receptor-ligand cargo complexes sorted by early endosomes is ligand-bound Tfn Receptor (TfR) (reviewed by (Antonescu et al., 2014)). Fe³⁺-bound Tfn (holo-Tfn) is taken up into cells through clathrin-mediated endocytosis while bound to TfR. Upon fusion of internalized vesicles with EEs, the acidified EE lumen triggers the release of iron from Tf (Klausner et al., 1983; Dautry Varsat et al., 1983). Fe³⁺ is subsequently reduced to Fe²⁺, while apo-Tfn remains bound to TfR, undergoing recycling to the plasma membrane. While Fe^2+^ can be transported into the cytosol via Divalent Metal Transporter 1 (DMT1) (Dev and Babitt, 2017), a portion of this labile Fe²⁺ pool is thought to be trafficked to mitochondria, yet the mechanisms facilitating its delivery remain incompletely defined. Interestingly, recent studies have revealed that EEs also form transient, non-fusogenic contacts with mitochondria, called “kiss-and-run” interactions, that may serve as localized sites of metabolite exchange (Hamdi et al., 2016; Barra et al., 2024; Das et al., 2016). The transfer of iron at these EE-mitochondria contact sites may also involve DMT1 (Barra et al., 2024).

The mechanisms that facilitate iron delivery to mitochondria may be particularly important when considering the metabolic need for iron in these organelles (Ben Zichri-David et al., 2025). Mitochondria are the primary consumers of intracellular iron, as mitochondrial iron is required for the synthesis iron–sulfur (Fe-S) clusters that drive electron transport and oxidative phosphorylation, as well as other biosynthetic reactions such as the production of heme (Ben Zichri-David et al., 2025). With the central role that iron metabolism plays in cell metabolism through the synthesis of Fe-S clusters, an important question that emerges is how iron delivery to mitochondria (e.g. from endosomes) is regulated to meet cellular requirements, particularly during conditions that trigger mitochondrial biogenesis and expansion.

Mitochondrial biogenesis occurs in response to several cues, including metabolic stress and nutrient insufficiency. AMP-activated protein kinase (AMPK) is a central metabolic sensor that becomes activated in response to a reduction of the ATP/AMP ratio or reduced glycolytic flux (Lin and Hardie, 2018; Herzig and Shaw, 2017; Trefts and Shaw, 2021; Steinberg and Hardie, 2022). AMPK triggers a range of signals that collectively increase nutrient uptake leading to ATP production and suppress some cellular processes, such as certain biosynthetic pathways. AMPK promotes mitochondrial biogenesis and enhances oxidative metabolism by several mechanisms (Herzig and Shaw, 2018), including phosphorylation of FNIP1 that promotes nuclear localization of TFEB (Malik et al., 2023). Importantly, mitochondrial biogenesis and expansion is expected to require an influx of iron so that oxidative capacity can scale with mitochondrial mass (Ward and Cloonan, 2018; Ben Zichri-David et al., 2025). However, whether and how AMPK may trigger increased mitochondrial iron influx has not been examined.

We previously showed that AMPK regulates endomembrane traffic, including the cell surface abundance and internalization of specific membrane proteins such as β1 integrin, implicating AMPK in regulation of endomembrane traffic at the level of plasma membrane (Ross et al., 2015, Orofiamma et al., 2025). However, whether AMPK also acts downstream of internalization to spatially reorganize endosomal compartments and facilitate inter-organelle communication has not been explored. Notably, the Rab5 and Rab31 guanine nucleotide exchange factor (GEF) Gapex-5 was identified as a novel AMPK substrate in a screen of novel targets (Ducommun et al., 2019). Gapex-5 regulates the internalization or sorting of specific receptors such as low-density lipoprotein receptor (LDLR) (Ducommun et al., 2019) and the glucose transporter GLUT4 (Lodhi et al., 2007). However, there are several GEFs that act on Rab5, including Rabex-5 (Yuan and Song, 2020), making the specific role of Gapex-5 in the regulation of Rab5-dependent sorting at EEs unclear. Moreover, the phosphorylation of Gapex-5 by AMPK suggests that AMPK may control early endosome dynamics by this mechanism, but the outcome of Gapex-5 phosphorylation by AMPK is poorly understood.

We examine the impact of AMPK activation on the spatial position of early endosomes, and the impact of this regulation on iron delivery to mitochondria. This work reveals that AMPK activation leads to the selective perinuclear clustering of early endosomes demarked by EEA1, without altering lysosome positioning. Using knockdown-rescue approaches with a S902A mutant of Gapex-5, we identify this AMPK phosphorylation site as critical for the AMPK-triggered perinuclear movement of EEs. We find that AMPK activation enhances the proximity of EEs with mitochondria in Gapex-5 dependent manner. We also reveal that AMPK triggers enhanced mitochondrial iron accumulation from endosomes in a Gapex-5 dependent manner. Together, these findings uncover a mechanism by which AMPK exerts control over endosome position via Gapex-5 phosphorylation and also triggers enhanced iron delivery to mitochondria.

## Results

### AMPK activation triggers perinuclear movement of early endosomes

To study the effects of AMPK activation on early endosome localization, we treated ARPE-19 cells with the AMPK agonist A-769662 for 15 min. While resting cells had punctate EEA1 staining consistent with distribution of EEA1 early endosomes throughout the cytoplasm, cells treated with A-769662 exhibited EEA1 punctate staining that was dramatically concentrated in the perinuclear region (**Figure 1A**). The distribution of EEA1 puncta was analyzed by kurtosis, which in this context measures the extent of fluorescence signal concentrated within a portion of the region of interest (Bone et al., 2017). In this application, an increase in kurtosis values reflect a concentration of EEA1 signal within a portion of the cell such as the perinuclear region. A-769662 treatment significantly increased the mean kurtosis of EEA1-positive early endosomes (**Figure 1B**), suggesting that AMPK activation altered the spatial distribution of EEs, leading to clustering of these organelles near the nucleus. To determine if this phenomenon was also observed when AMPK activation occurred in response to metabolic stress, we examined treatment with 2-deoxyglucose (2DG) to inhibit glycolysis or oligomycin to inhibit mitochondrial ATP synthesis. We previously showed that these treatments trigger AMPK activation in ARPE-19 cells (Orofiamma et al., 2025). Both 2DG and oligomycin treatments triggered the redistribution of EEA1 early endosomes to the perinuclear region (**Figure 1C-F**).

**Figure 1.**
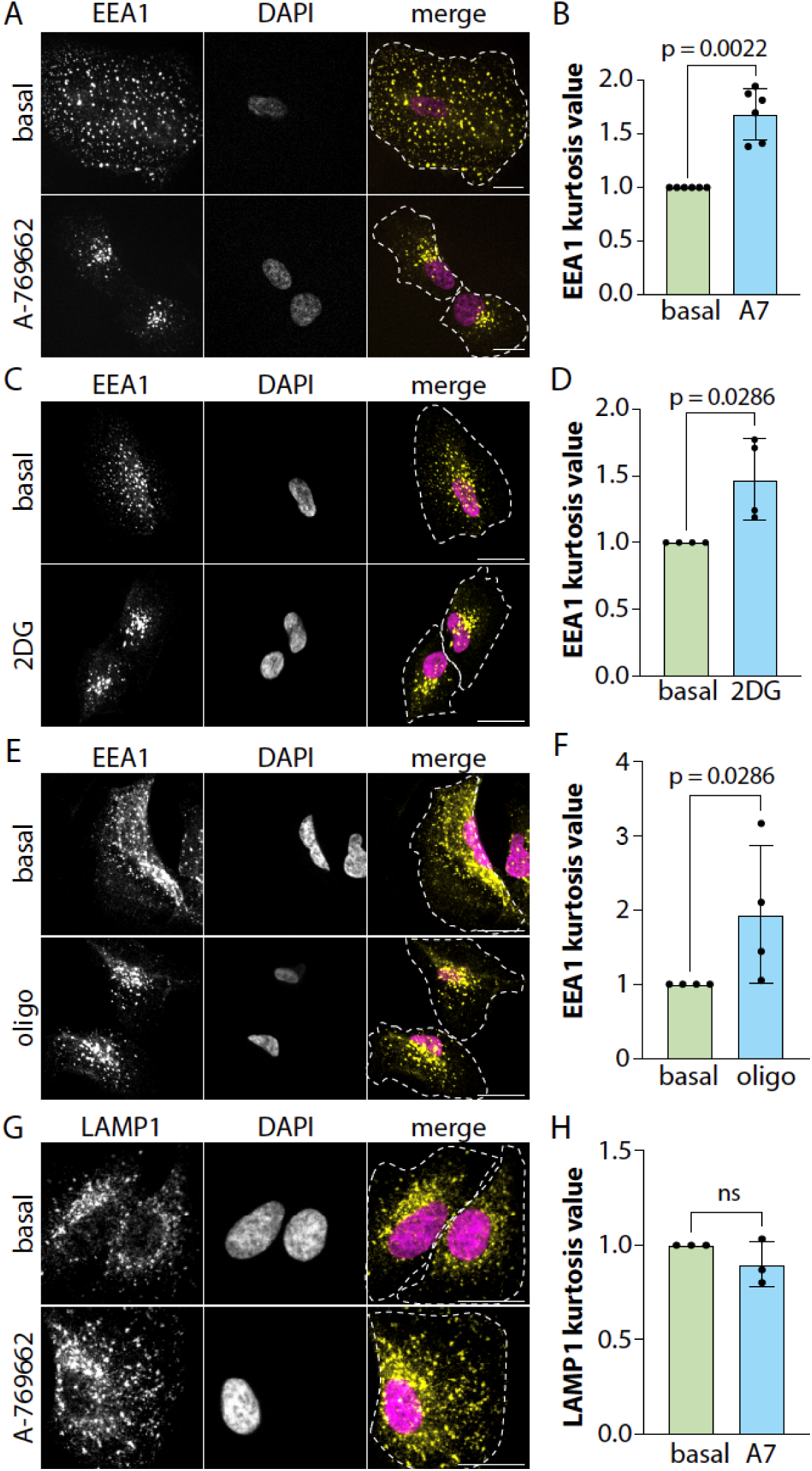
AMPK activation triggers the selective perinuclear repositioning of early endosomes. ARPE-19 cells were treated with 100 μm A769662 or vehicle (0.1% v/v DMSO) control for 15 min (***A-B,G-H***), 25 mM 2-deoxyglucose or vehicle (media) for 15 min (***C-D***), or 5 mM oligomycin or vehicle control for 20 min (***E-F***). Shown for each are representative images obtained by spinning disc confocal microscopy, scale 20 μm (A, C, E, G) and quantification of kurtosis value, shown as mean ± SD from 6 (B), 4 (D), 4 (F), or 3 (H) independent experiments, with each experiment quantifying at least 30 cells per condition (normalized to basal condition). All statistical comparisons were performed by Mann-Whitney test.

The redistribution of early endosomes to the perinuclear region upon AMPK activation could reflect the specific regulation of this organelle or broader control of membrane compartments. A-769662 treatment was without effect on the distribution of lysosomes detected by LAMP1 staining (**Figure 1G-H**). AMPK activation also leads to inhibition of mTORC1 (Smiles et al., 2024), so we next examined whether early endosome redistribution is specific to AMPK activation or can also be induced by mTORC1 inhibition. Cells treated with the mTORC1 inhibitor Torin exhibited no significant differences in the spatial position of EEA1 early endosomes (**Figure S1**). This suggests that suppression of mTORC1 signaling does not induce the spatial redistribution of early endosomes. Collectively, these results indicate that AMPK activation triggers the selective perinuclear redistribution of early endosomes.

### Gapex-5 is required for AMPK-stimulated perinuclear localization of early endosomes

We next probed the role of the AMPK substrate Gapex-5 in AMPK-mediated early endosomal perinuclear redistribution (**Figure 2A**). To do so, we established effective siRNA silencing of Gapex-5 expression (**Figure S2A**). While cells treated with non-targeting siRNA exhibited a significant increase in perinuclear clustering upon treatment with A-769662, this treatment was without appreciable effect in cells subjected to Gapex-5 silencing (**Figure 2B**).

**Figure 2.**
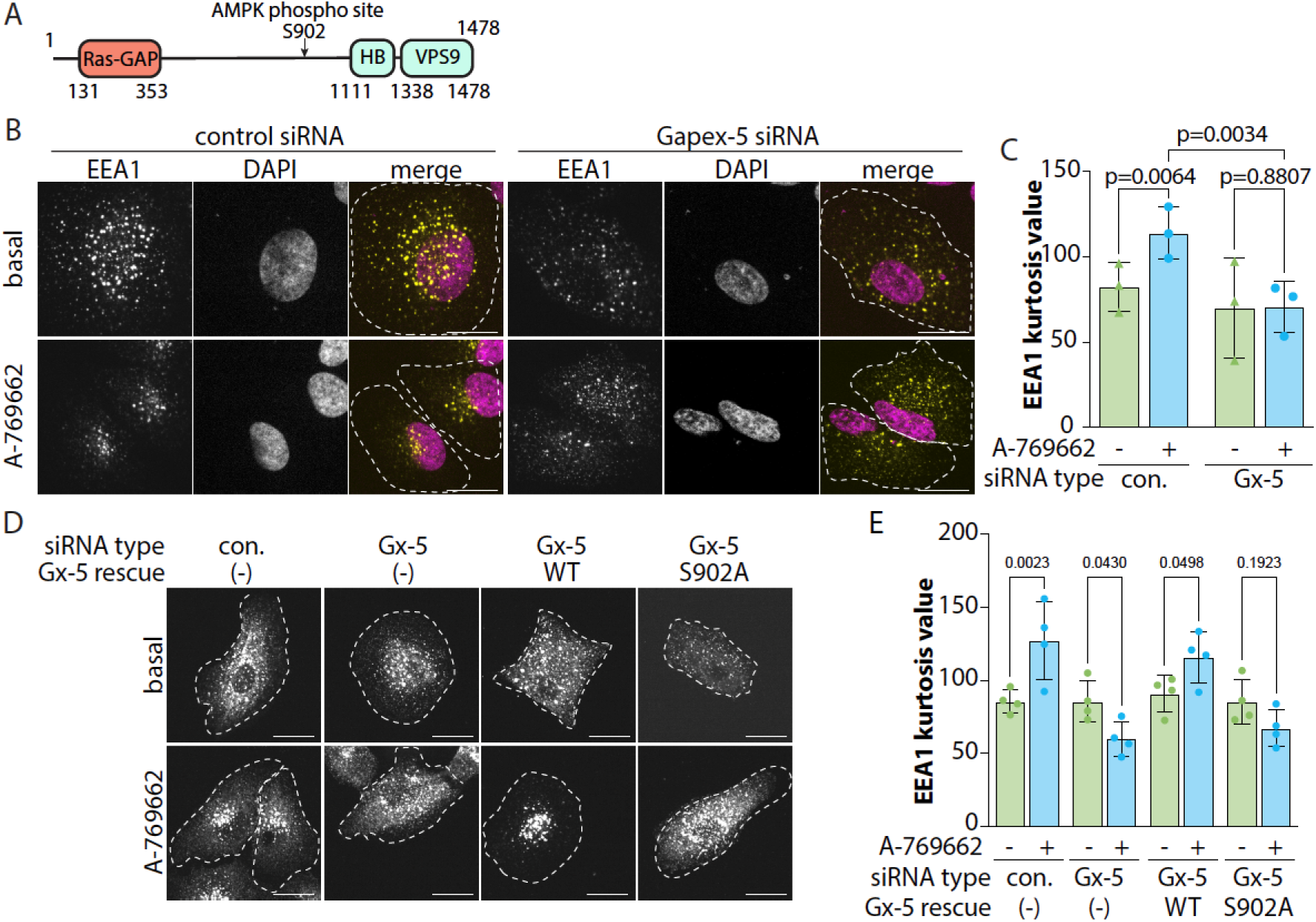
Early endosome perinuclear movement triggered by AMPK activation requires Gapex-5. (A) Diagram of Gapex5 showing Ras-GAP, HB, and Vps9 domains, as well as the previously identified AMPK phosphorylation site. (B-C) ARPE-19 cells were subjected to siRNA silencing using sequences targeting Gapex-5 (Gx-5) or non-targeting control (con.). Cells were treated with 100 μm A769662 or vehicle (0.1% v/v DMSO) control for 15 min. Shown in (B) are representative images obtained by spinning disc confocal microscopy, scale 20 μm and in (C) quantification of kurtosis value, shown as mean ± SD from 3 independent experiments, with each experiment involving measurement of at least 30 cells per condition. Statistical comparisons were performed by 2-way ANOVA with Fisher’s Least Significant Difference (LSD). (D-E) ARPE-19 cells engineered using the Sleeping Beauty transposon system to carry a stable transgene for doxycycline-inducible expression of eGFP-Gapex5 WT or S902A were subjected to siRNA silencing using sequences targeting Gapex-5 (Gx-5) or non-targeting control (con.). Expression of eGFP-Gapex5 constructs was induced with 0.5 μM doxycycline for 24h (rescue conditions), or left uninduced, depicted by (-). Shown in (D) are representative images obtained by spinning disc confocal microscopy, scale 20 μm and in (E) quantification of kurtosis value, shown as mean ± SD from 4 independent experiments, with each experiment involving measurement of at least 30 cells per condition. Statistical comparisons were performed by 2-way ANOVA with Šidák post-hoc test.

To directly test the role of S902 phosphorylation of Gapex-5 by AMPK, we established a knockdown-rescue approach. We generated ARPE-19 cell lines that stably carry a transgene that allows doxycycline inducible expression of either wild-type (WT) or a phosphorylation deficient mutant (S902A) Gapex-5 fused to eGFP using the Sleeping Beauty transposon system (Kowarz et al., 2015), as we have done previously (Zak and Antonescu, 2023; Cabral-Dias et al., 2022; Orofiamma et al., 2025; Rahmani et al., 2023) (**Figure S2B**). Cells subjected to Gapex-5 silencing in the absence of rescue by doxycycline treatment exhibited a suppression of the perinuclear movement of early endosomes upon treatment with A-769662 (**Figure 2C**). In contrast, rescue of Gapex-5 silencing by expression of WT eGFP-Gapex5 restored the perinuclear movement of EEA1 early endosomes in cells treated with A-769662. Importantly, similar expression of eGFP-Gapex-5 S902A failed to rescue the effect of Gapex-5 silencing on EEA1 early endosomes upon A-769662 treatment. These results indicate that AMPK activation requires Gapex-5, and specifically the S902 site on Gapex-5 previously shown to be phosphorylated by AMPK (Ducommun et al., 2019), for redistribution of early endosomes to the perinuclear region.

### Early endosome repositioning upon AMPK activation requires microtubules and the dynein adaptor Hook3

Early endosome movement within cells is coordinated by microtubule motors. Hook3 is part of the FHF complex that acts as a dynein adaptor, allowing minus-end movement of early endosomes along microtubules. We used silencing of Hook3 (**Figure S3**) to assess its role in the perinuclear early endosome repositioning driven by AMPK activation. In cells treated with non-targeting siRNA, A-769662 treatment triggered a perinuclear clustering of EEA1 early endosomes (**Figure 3A-B**). In contrast, in cells treated with Hook3 siRNA, EEA1 early endosomes remained distributed throughout the cell upon A-769662 treatment (**Figure 3A-B**). Consistent with this, treatment of cells with nocodazole to disrupt microtubules blunted the effect of A-769662 treatment on the perinuclear redistribution of EEA1 early endosomes (**Figure 3C-D**).

**Figure 3.**
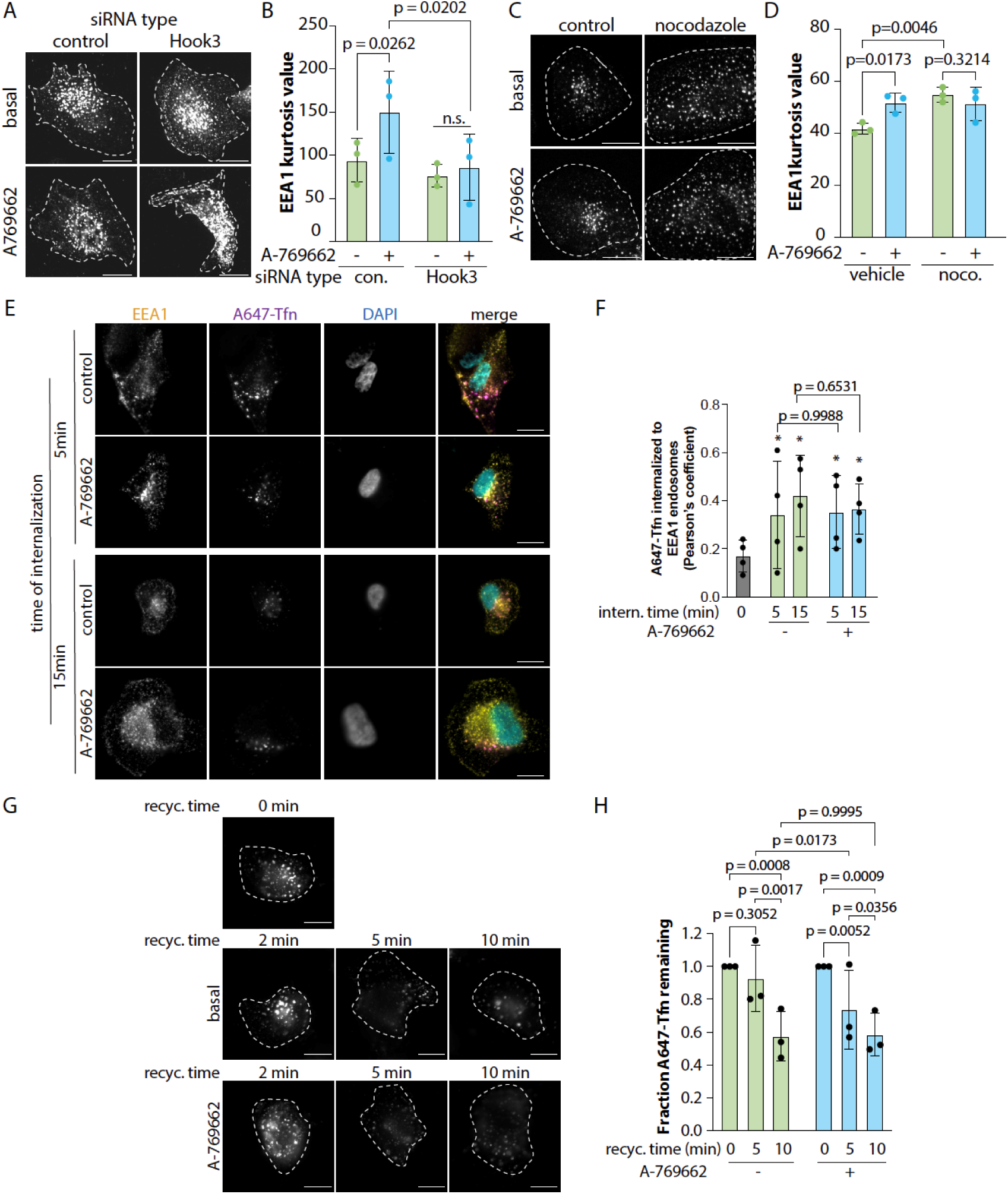
Early endosome perinuclear movement triggered by AMPK activation requires Hook3 yet minimally impacts transferrin receptor endosomal sorting. (***A-B***) ARPE-19 cells were subjected to siRNA silencing using sequences targeting Hook3 or non-targeting control (con.). Cells were treated with 100 μm A769662 or vehicle (0.1% v/v DMSO) control for 15 min. Shown in (A) are representative images obtained by spinning disc confocal microscopy, scale 20 μm and in (B) quantification of kurtosis value, shown as mean ± SD from 3 independent experiments, with each experiment involving measurement of at least 30 cells per condition. (***C-D***) ARPE-19 cells were subjected to treatment with nocodazole (noco.) or vehicle control for 24h. Cells were then treated with 100 μm A-769662 or vehicle (0.1% v/v DMSO) control for 15 min. Shown in (C) are representative micrographs obtained by spinning disc confocal microscopy, scale 20 μm and in (D) quantification of kurtosis value, shown as mean ± SD from 3 independent experiments, with each experiment involving measurement of at least 30 cells per condition. (***E-F***) ARPE-19 cells were treated with 100 μm A769662 or vehicle (0.1% v/v DMSO) control for 15 min, followed by A647-Tfn internalization assay. Representative fluorescence micrographs obtained by spinning disc confocal microscopy (E), and the extent of colocalization determined by Pearson’s coefficient, shown as mean ± SD from 4 independent experiments, with each experiment involving measurement of at least 30 cells per condition. (***G-H***) ARPE-19 cells were treated with 100 μm A769662 or vehicle (0.1% v/v DMSO) control for 15 min, followed by A647-Tfn recycling assay. Representative fluorescence micrographs obtained by spinning disc confocal microscopy (E), and the amount of A647-Tfn remaining in cells determined by mean fluorescence intensity, shown as mean ± SD from 3 independent experiments, with each experiment involving measurement of at least 30 cells per condition. All statistical comparisons were performed by 2-way ANOVA with Tukey post-hoc test.

The perinuclear movement of early endosomes along microtubules could impact the arrival of holo-Tfn in EEA1 early endosomes upon its internalization from the cell surface by clathrin-mediated endocytosis. To examine this, we performed internalization experiments using fluorescently labeled holo-Tfn (A647-Tfn). Following 5 min or 15 min of A647-Tfn internalization, we observed partial co-localization with EEA1, consistent with normal early endosomal trafficking kinetics (**Figure 3E**). Pearson’s coefficient of A647-Tfn and EEA1 endosomes at 5-15 min was increased compared to t=0 min of A647-Tfn internalization (addition of labeled ligand followed by its near-immediate washout) (**Figure 3F**). Treatment with A-769662 did not appreciably alter Tfn arrival to EEA1 endosomes (**Figure 3E-F**). We also examined Tfn recycling by labeling cells with A647-Tfn for 15 min, followed by washing to remove unbound A647-Tfn and then incubating cells in media lacking A647-Tfn. In this assay, the loss of A647-Tfn from cells measures the rate of TfR recycling to the plasma membrane from intracellular compartments. A-769662 treatment led to a minor enhancement of the loss of A647-Tfn at 5 min of recycling, while this treatment had no effect at 10 min of recycling. These data indicate that AMPK activation does not have a substantial effect on the normal routing of TfR to early endosomes or its recycling to the plasma membrane following internalization.

### AMPK activation increases early endosome proximity to mitochondria in a Gapex-5 dependent manner

The redistribution of EEA1 early endosomes to the perinuclear region implies that the proximity of these organelles relative to other compartments could support specific functions during metabolic stress. Mitochondria exhibit significant perinuclear localization in many cells, including in ARPE-19 cells. To resolve whether AMPK-triggered EEA1 early endosome perinuclear clustering could potentially result in an alteration of early endosome proximity to mitochondria, we examined the co-localization of EEA1-labeled early endosomes with mitochondria labeled with MitoTracker by spinning disc confocal microscopy. We performed this experiment in conjunction with siRNA silencing of Gapex-5. In cells treated with non-targeting (control) siRNA, we observed that A-769662 treatment elicited an increase in the co-localization of EEA1 early endosomes and mitochondria compared to basal conditions (**Figure 4A-B**). In contrast, there was no appreciable change in the overlap of EEA1 early endosomes and mitochondria upon A-769662 treatment in Gapex-5 silenced cells (**Figure 4A-B**). This indicates that AMPK activation may increase the proximity of early endosomes to mitochondria, and that this occurs in a Gapex-5 dependent manner.

**Figure 4.**
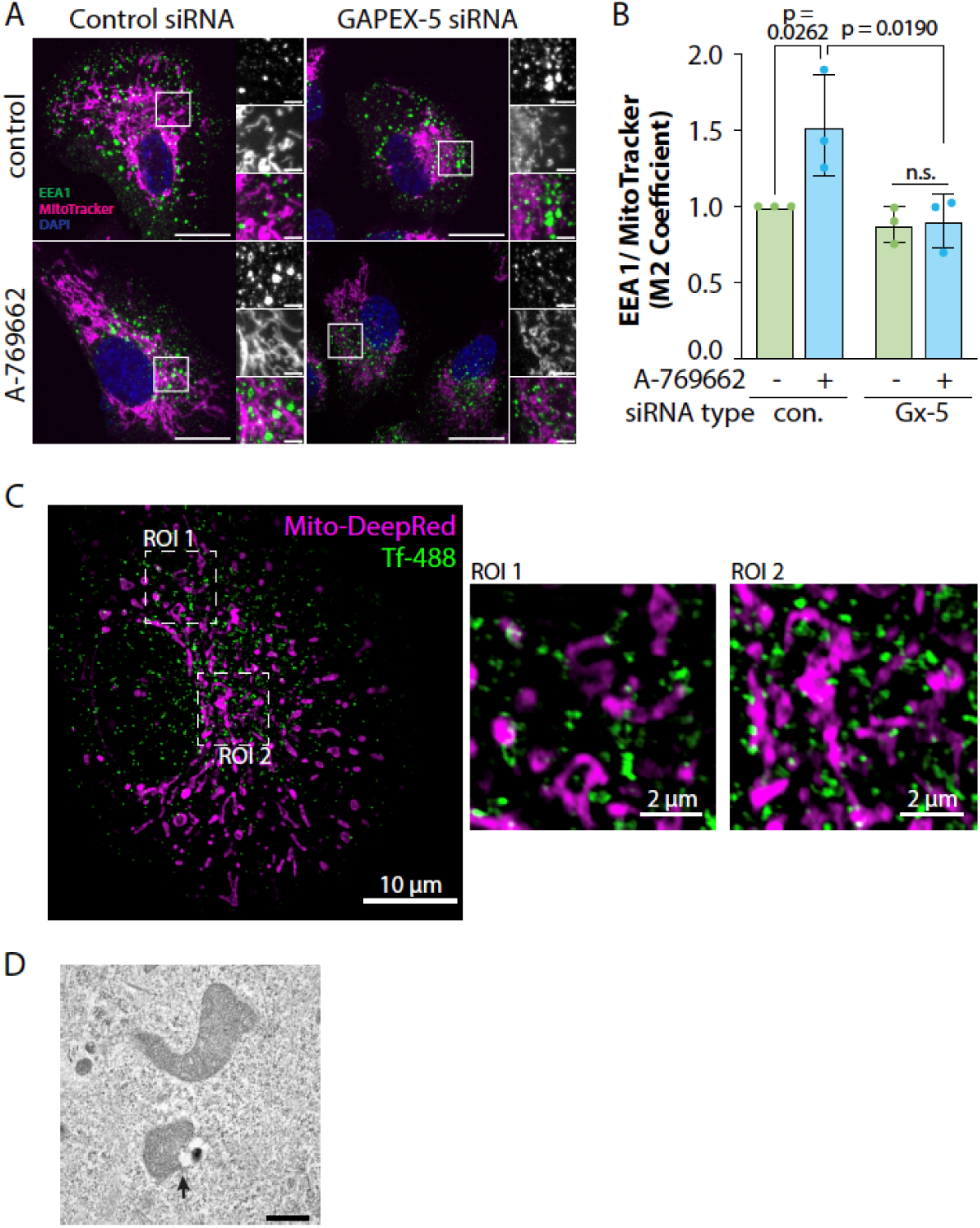
AMPK activation triggers enhanced proximity of early endosomes with mitochondria. (***A-B***) ARPE-19 cells were treated with 100 μm A769662 or vehicle (0.1% v/v DMSO) control for 15 min, followed by treatment with MitoTracker Deep Red and antibody labeling targeting EEA1. Shown in (A) are representative images obtained by spinning disc confocal microscopy, scale 20 μm and in (B) quantification of co-localization between EEA1 and MitoTracker by Manders’ coefficient, shown as mean ± SD from 3 independent experiments (normalized to control siRNA, untreated), with each experiment involving measurement of at least 30 cells per condition. Statistical comparisons were performed by 2-way ANOVA with Fisher’s Least Significant Difference (LSD). (C) ARPE-19 cells were incubated with Alexa488 conjugated human transferrin for 1h and MitoTracker Deep Red for 20 min, followed by fixation and imaging using SIM (D) ARPE19 cells were treated with 100 μm A769662 for 15 min, and then were fixed in 4% paraformaldehyde and 0.1% glutaraldehyde in PBS, followed by sample preparation and transmission electron microscopy as in (Hua et al., 2015), scale 500 nm.

Lysosomes also exhibit some perinuclear concentration in many cells, including ARPE-19 cells. To determine if the increased overlap of EEA1 early endosomes with mitochondria reflected an increased concentration of endosomes and mitochondria in a crowded perinuclear region, we examined the overlap of EEA1 endosomes with LAMP1 lysosomes. We observed no changes in the overlap of EEA1 early endosomes and LAMP1 lysosomes upon treatment with A-769662 (**Figure S4**), suggesting that the AMPK-directed enhanced proximity of EEA1 with mitochondria reflects a specific association and not solely crowding of organelles in the perinuclear region.

To obtain better insight into the position of early endosomes relative to mitochondria upon AMPK activation, we performed structured illumination microscopy (**Figure 4C**) and transmission electron microscopy (**Figure 4D**). In response to A-769662 treatment, we observed that early endosomes labeled with fluorescent transferrin internalized from the cell surface formed many very close contacts with mitochondria (**Figure 4C**), which is consistent with the close proximity of mitochondria and membrane-bound structures consistent with endosomes observed by transmission electron microscopy (**Figure 4D**). Together with the observations of AMPK-stimulated increased in proximity of EEA1 early endosomes to mitochondria by spinning disc confocal microscopy (**Figure 4A-B**), these results are consistent with AMPK-stimulated redistribution of early endosomes to the perinuclear region, supporting contacts of early endosomes with mitochondria.

### AMPK activation triggers increased delivery of endosomal iron to mitochondria

One of the actions of AMPK is to promote ATP generation under conditions of nutrient stress. To examine this, we measured mitochondrial membrane potential using TMRE, a mitochondrial membrane potential sensitive fluorescent dye. Treatment with A-769662 elicited an increase in TMRE fluorescence, indicative of an increase in mitochondrial membrane potential (**Figure S5A-B**). This is consistent with an increase in mitochondrial metabolism, which could reflect an increase in central carbon metabolism flux, or an expansion of mitochondrial enzymes, or both. Importantly, many of the enzymes involved in the TCA cycle and the electron transport chain require iron in the form of Iron-Sulphur (Fe-S) clusters (Ben Zichri-David et al., 2025). This suggests that under conditions of nutrient stress that lead to AMPK activation, mitochondria may have an increased demand for iron.

The increased proximity of EEA1 early endosomes with mitochondria (**Figure 4**), consistent with enhanced membrane contacts between these organelles upon AMPK activation. This suggests that AMPK may promote molecular exchange between these organelles dependent on endosome-mitochondria membrane contacts. To test this, we next examined whether AMPK activation triggers an increase in iron delivery to mitochondria using the mitochondrial iron fluorescent probe mito-FerroGreen (MFG) (Hirayama et al., 2018). We observed that treatment with A-769662 for 4h resulted in a significant increase in MFG signal (**Figure 5A-B**). Importantly, this MFG signal was virtually eliminated upon treatment with the iron chelator deferoxamine, indicating that the signal reflects iron-dependent fluorescence. (**Figure 5A-B**).

**Figure 5.**
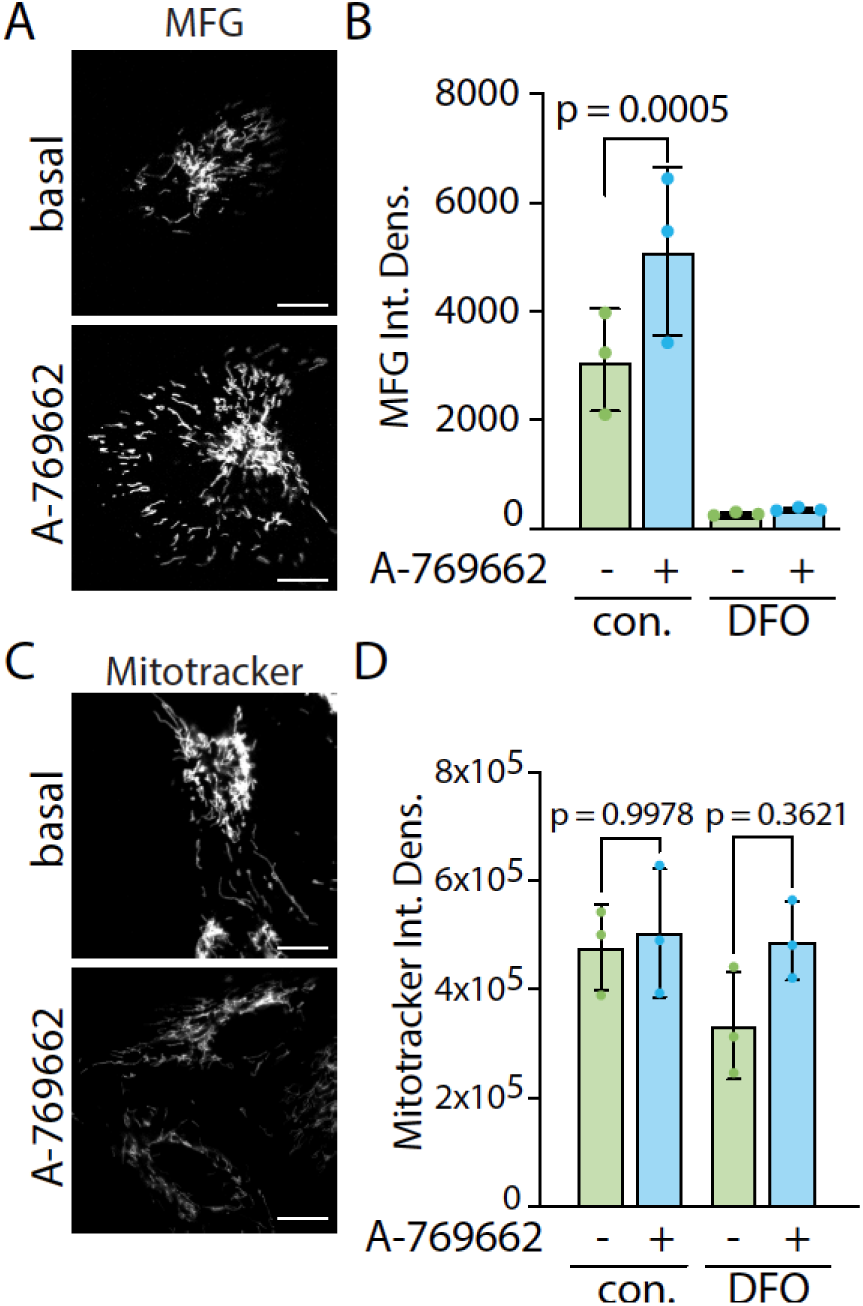
AMPK activation triggers an increase in mitochondrial iron. ARPE-19 cells were treated with 100 μm A769662 or vehicle (0.1% v/v DMSO) control for 15 min; samples as indicated were also treated with deferoxamine (DFO, 10 mM for 1 h, followed by treatment labeling with MitoFerroGreen (MFG) (A-B) or MitoTracker Deep Red (C-D) followed by live-cell spinning disc confocal microscopy. Shown in (A, C) are representative micrographs obtained, scale 20 μm and in (B,D) quantification of integrated fluorescence intensity, shown as mean ± SD from 3 independent experiments, with each experiment involving measurement of at least 30 cells per condition.

AMPK activation could trigger an increase in mitochondrial mass rather than a selective increase in mitochondrial iron. However, similar treatment with A-769662 for 4h did not change the total intensity of MitoTracker staining (**Figure 5C-D**). Consistent with known effects of AMPK to promote mitochondrial fission (Toyama et al., 2016), we observed that A-769662 treatment for 4h led to an increase in mitochondrial fragmentation (**Figure S5C**). Together, these results indicate that AMPK activation triggers an increase in mitochondrial iron that cannot be accounted for by an increase in mitochondrial mass.

The increase in mitochondrial iron upon AMPK activation could be derived from cytosolic stores of iron, or from delivery of iron from endosomes. To probe this, we suppressed early endosomal traffic by treatment with VPS34-IN1, an inhibitor of the class III phosphoinositide-3-kinase Vps34 that produces phosphatidylinositol-3-phosphate essential for membrane traffic and sorting at the early endosome (Wang et al., 2019; Gautreau et al., 2014). Treatment with VPS34-IN1 abolished the increase in mitochondrial iron detected by MFG observed upon A-769662 treatment (**Figure 6A-B**).

**Figure 6.**
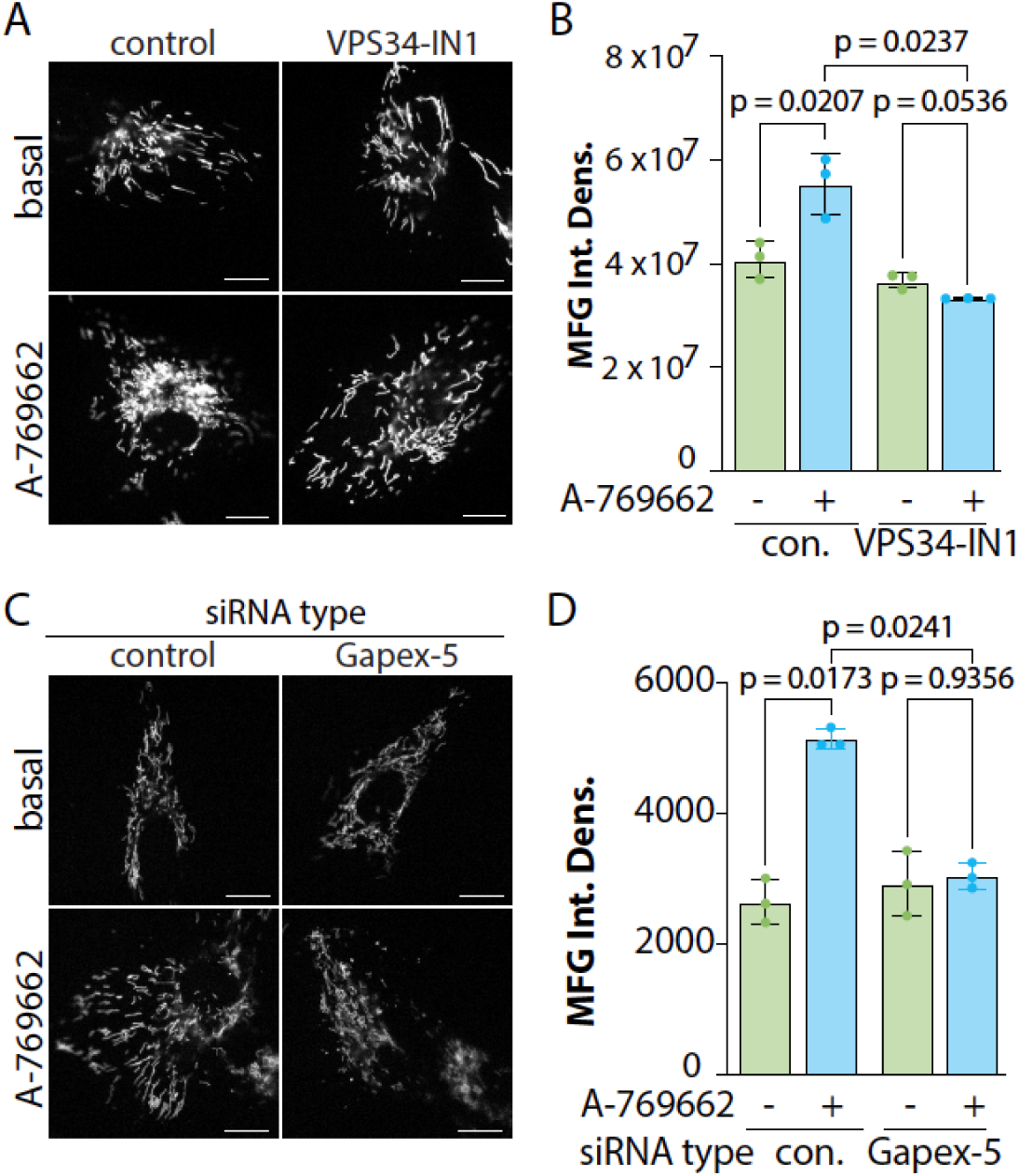
The increase in mitochondrial iron upon AMPK activation requires endosomal phosphatidylinositol-phosphate and Gapex-5. ARPE-19 cells were either (***A-B***) treated with 1 μM VPS34-IN1 or vehicle control (DMSO, 0.1% v/v), or (***C-D***) subjected to siRNA silencing using sequences targeting Gapex-5 or non-targeting control (con.). Cells were treated with 100 μm A769662 or vehicle (0.1% v/v DMSO) control for 15 min, followed by treatment labeling with MitoFerroGreen (MFG) followed by live-cell spinning disc confocal microscopy. Shown in (A, C) are representative micrographs obtained, scale 20 μm and in (B,D) quantification of mean fluorescence intensity, shown as mean ± SD from 3 independent experiments, with each experiment involving measurement of at least 30 cells per condition. All statistical comparisons were performed by 2-way ANOVA with Tukey post-hoc test.

We next examined the role of Gapex-5 in mitochondrial iron delivery. While cells treated with non-targeting siRNA exhibited an increase in MFG signal upon treatment with A-769662, Gapex-5 silenced cells exhibited no change in MFG signal upon A-769662 treatment (**Figure 6E-F**). Together, these results indicate that AMPK activation triggers an increase in endosomal delivery of iron to mitochondria in a Gapex-5 dependent manner.

## Discussion

### Distinct metabolic signals regulate movement and position of endosomes and lysosomes

Our observation that the movement and positioning of early endosomes are regulated by nutrient cues via AMPK is consistent with evidence that other endocytic compartments, particularly lysosomes, are also subject to metabolic control. Lysosome function and lumenal pH are regulated by their subcellular location, with peripheral lysosomes being less acidic than perinuclear ones (Johnson et al., 2016). The spatial arrangement of lysosomes is gated by sensing of amino acids by the BORC complex, which controls the recruitment of the microtubule plus-end directed motors Kif1B and Kif5B to lysosomes via the adaptor protein Arl8b (Pu et al., 2015, 2017). In the presence of amino acids, the peripheral movement of lysosomes by this system is critical to allow a range of functions including Akt signaling (Jia and Bonifacino, 2019). This establishes a model in which lysosome movement is regulated by changes in the recruitment of microtubule motor protein adaptors such as Arl8b to that organelle.

The position of early endosomes is regulated by specific motor protein adaptors and regulators that are at least in part distinct from those that govern lysosome position. Early endosome movement is regulated by FHF complexes. FHF complexes containing Hook3 are autoinhibited, with relief of the autoinhibition of Hook3 occurring upon binding of the kinesin Kif1C to Hook3, which allows Hook3 to engage with dynein-dynactin and support minus-end directed microtubule movement (Abid Ali et al., 2025). As plus-end directed microtubule movement of early endosomes is promoted by Kif1C, the relative impact of Kif1C-dependent vs FHF/Hook3-dependent early endosome movement may underlie the ability of early endosomes to undergo perinuclear vs peripheral movement. Interestingly FHF complexes containing Hook1 or Hook3 interacted selectively with Rab5-GTP but not with Rab5-GDP and regulated the position of TfR containing endosomes in neurons (Guo et al., 2016). This suggests that modulation of Rab5-GTP loading upon AMPK activation, perhaps resulting from phosphorylation of Gapex-5, could modulate Kif1C-*vs* FHF/Hook3-dependent endosomal movement. Resolving the mechanism by which AMPK mediates this regulation is beyond the scope of this study and insights into this from future work will be very informative.

While we establish that S902 on Gapex-5 is required for the AMPK-driven perinuclear movement of early endosomes and endosomal iron delivery to mitochondria, how AMPK phosphorylation of Gapex-5 on S902 regulates Gapex-5 activity and function is not clear. One possibility is that phosphorylation of Gapex-5 on S902 increases its GEF activity towards Rab5 (Guo et al., 2016), allowing increased Rab5 GTP binding on early endosomes that may enhance interaction with effectors such as FHF complexes that contain Hook3 (Guo et al., 2016), thus promoting perinuclear movement. AMPK enhances Rab5 GTP loading (Rao et al., 2021) but may also suppress Rab5 levels within minutes (Diesendorf et al., 2025). There are many Rab5 effectors, and interaction with specific effectors appears to be context-specific, as Rab5 effectors EEA1 and APPL1 are recruited to different subsets of endosomes (Kalaidzidis et al., 2015; van der Beek et al., 2022). In this light, it is also possible that S902 phosphorylation of Gapex-5 does not control GEF activity, but instead could impact possible scaffolding functions by modulating interactions with other proteins. As such, while it is beyond the scope of this current study, future studies that may reveal the outcome of Gapex-5 phosphorylation by AMPK on its GEF activity and other functions will provide very useful information.

### Regulation of mitochondrial iron and metabolism by AMPK

We uncover a novel mechanism for the regulation of mitochondrial iron following AMPK activation that requires Gapex-5 and that correlates with AMPK-stimulated movement of early endosomes to the perinuclear region. Mitochondria are the major destination of iron in a cell, and mitochondrial iron is used for biosynthesis of Fe-S clusters and heme (Ben Zichri-David et al., 2025). In turn, both Fe-S clusters and heme function as cofactors for a range of enzymes. Estimates suggest that hundreds of human enzymes bind iron, a significant portion of which is in the form of Fe-S clusters. In the mitochondria, Fe-S clusters are essential cofactors for enzymes in the TCA cycle in which aconitase, which catalyzes the conversion of citrate to isocitrate, binds an Fe-S cluster in its active site (Read et al., 2021). Fe-S clusters are also important in the electron transport chain; for example complex I binds up to 10 Fe-S clusters (Gnandt et al., 2016). As a result, the availability of iron delivery to mitochondria is expected to be a limiting factor for enzymes that depend on Fe-S clusters, in particular the enzymes required for mitochondrial ATP production.

Iron homeostasis is important and must be tightly regulated, as both iron deficiency and iron overload can lead to mitochondrial damage (Walter et al., 2002). The expression of mitochondrial ferritin (MtFt) allows mitochondria to store and sequester a supply of iron that is approximately 30-50% of the iron storage of a cell (Levi et al., 2021; Jhurry et al., 2012). Importantly, there is evidence to support the model in which mitochondrial iron is limiting for ATP production and mitochondrial metabolism. For example, increased expression of MtFt leads to a shift in iron from the cytosol to the mitochondria, which is associated with an increase in Fe-S dependent mitochondrial enzyme activity and ATP levels (Campanella et al., 2008). However, a separate study found that MtFt overexpression could lead to sequestration of iron in the mitochondria, rendering it less available for Fe-S dependent mitochondrial enzymes (Nie et al., 2005). While there are likely context-specific outcomes to MtFt function, these studies are consistent with the activity of Fe-S dependent mitochondrial enzymes being dependent on iron availability in mitochondria. As such, an AMPK-stimulated increase in free mitochondrial Fe^2+^ as we have observed here is consistent with an increase in mitochondrial respiration and ATP production driven by AMPK activation.

Previous studies identified close contacts between mitochondria and early endosomes, suggesting that these may allow direct transfer of iron to mitochondria (Hamdi et al., 2016; Barra et al., 2024; Das et al., 2016), and implicate the transporter Dmt1 in this phenomenon (Barra et al., 2024). Our work identifies that AMPK regulates this phenomenon, and the signals and mechanism by which this regulation takes place. Interestingly, the mitochondrial fusion regulator mitofusin (Mfn1/2) interacts with Rab5c (Irazoki et al., 2023), allowing tethering and close apposition of these organelles. The molecular mechanisms underlying possible membrane contact sites between endosomes and mitochondria remain to be elucidated.

### Metabolic regulation of endomembrane network sorting

We identified that AMPK regulates the position of early endosomes in a Gapex-5 dependent manner. Interesting, AMPK activation had no discernable impact on the arrival of transferrin into early endosomes following its internalization, and only a modest effect on its recycling back to the plasma membrane. This is consistent with previous studies that found that AMPK activation did not alter Tfn internalization or cell surface TfR levels (Ross et al., 2015; Orofiamma et al., 2025). This suggests that even though EEA1 early endosomes transit away from the cell periphery towards the perinuclear area, Tfn-loaded vesicles internalized from the cell surface may also similarly undergo this perinuclear transit.

While the endomembrane traffic and sorting of TfR is not robustly regulated by AMPK activation, other cargo receptors are more profoundly impacted by these signals. We recently reported that the clathrin-mediated internalization of β1-integrin is significantly increased upon AMPK activation, as a result of engagement of a pathway involving Arf6 and Dab2 (Orofiamma et al., 2025). This adds to a growing appreciation that AMPK regulates the sorting of specific receptors and transporters along the endomembrane network, including the Na/K ATPase (Gusarova et al., 2009), glucose transporters (Wu et al., 2013; Habegger et al., 2012; Antonescu et al., 2008a; Yang and Holman, 2005), and CD36 (Luiken et al., 2003). In the context of AMPK regulation and phosphorylation of Gapex-5, it is notable that Gapex-5 perturbation caused disruption of internalization of the low-density lipoprotein receptor (LDLR) (Ducommun et al., 2019) or disruption of intracellular retention of GLUT4 glucose transporters (Lodhi et al., 2007). Hence, while AMPK activation did not lead to changes in the arrival of TfR to EEA1 early endosomes or its departure therefrom, the regulation of Gapex-5 by AMPK may alter the sorting of other cargo. Thus, in addition to providing insights into regulation of endosomal iron to mitochondria, this work may inform future studies that examine control of endomembrane traffic and sorting by AMPK.

In summary, we identified a novel action of AMPK activation that regulates the position of early endosomes, which may facilitate the delivery of endosomal iron to mitochondria. This adds to the growing appreciation of metabolic regulation of the endomembrane network and the important impact that this has on cellular adaptation to fluctuating nutrient conditions.

## Acknowledgements

This work was supported by Discovery Grants from the Natural Sciences and Engineering Research Council (NSERC) to C.N.A. (RGPIN-2016-04371).

## Materials and Methods

### Materials

Details of reagents and materials are provided in **Table S1**.

### Cell Culture

Human non-immortalized retinal pigment epithelial cells (ARPE-19) obtained from the American Type Culture Collection [ATCC, Manassas, VA]) were cultured in Dulbecco’s Modified Eagle’s Medium/Nutrient Mixture F-12 (DMEM/F-12, 1:1; Gibco, Thermo Fisher Scientific) supplemented with 10% fetal bovine serum (FBS; Thermo Fisher Scientific) and 100 U/mL penicillin and 100 μg/mL streptomycin (P/S; Gibco, Thermo Fisher Scientific). Cells were maintained at 37C in a humidified incubator with 5% CO₂. Cultures were passaged at ∼80% confluency using 0.25% Trypsin-EDTA (Gibco).

### Cell treatments and fluorescent probe labeling

All experiments began with cells incubated in no serum (0% FBS) culture media for 1 h before experimental treatments. Allosteric AMPK activation was induced with 100 μM A-769662 (Abcam) or an equal volume of DMSO (vehicle control; BioShop). Other pharmacological treatments were performed with 5 μM oligomycin (Cell Signaling Technology) or DMSO, 25 mM 2-deoxyglucose (2-DG; Millipore Sigma), with corresponding volume of water as a vehicle control. Class III phosphatidylinositol 3-kinase (PI3K) inhibition was achieved using 1 μM VPS34-IN1 (Cayman Chemical) or DMSO for 4 h. To manipulate iron availability, cells were treated with 100 μM deferoxamine mesylate (DFO) (Sigma Aldrich, D9533), an iron chelator, or 50 μg/mL holo-transferrin (Sigma Aldrich, T0665), for 4 h. For visualization of mitochondrial iron, MitoFerroGreen (MFG; 5 μM; Dojindo) was applied for 30 min prior to imaging, protected from light. MitoTracker™ Deep Red FM (200 nM; Thermo Fisher Scientific) was used in selected experiments to assess mitochondrial content and overlap with EEA1.

### Stable transfections using Sleeping Beauty transposon system

pSBtet-BP was a gift from Eric Kowarz (Goethe-University of Frankfurt, Frankfurt, Germany, plasmid 60496; Addgene; http://n2t.net/addgene:60496; RRID:Addgene_60496) (Kowarz et al., 2015). pCMV(CAT)T7-SB100 was a gift from Zsuzsanna Izsvak (Max Delbrück Center for Molecular Medicine, Berlin-Buch, Germany, plasmid 34879; http://n2t.net/addgene:34879; Addgene; RRID:Addgene_34879) (Mátés et al., 2009). To create the ARPE-19 cell lines that carry a stable transgene for inducible expression of mStayGold fused to Gapex-5 (WT), an oligonucleotide encoding Gapex-5 fused to eGFP was generated by Genscript using the ORF sequence of mStayGold, as per (Ivorra-Molla et al., 2023), followed by a spacer peptide sequence (5′-GGGGGGTCTGGTGGCAGTGGAGGGGGATCC-3′), followed by an oligonucleotide sequence encoding Gapex-5 as per GenBank accession number NM_001282679.2. This oligonucleotide sequence was subcloned into pSBtet-BP to generate pSBtet-BP-mSG-Gapex-5 WT. To create a corresponding plasmid encoding S902A Gapex-5, the codon encoding for S902 (AGC) was mutated to GCA (Alanine).

pSBtet-BP plasmids encoding mSG-Gapex5 WT or S902A were co-transfected with pCMV(CAT)T7-SB100 into ARPE-19 cells using FuGENE HD transfection reagent as per the manufacturer’s protocol (Promega). Selection of stably engineered cells was performed in growth media supplemented with 2 μg/mL puromycin for 2-3 weeks. To induce the expression of mSG-Gapex5 constructs, cells were incubated with 0.5 μM doxycycline (dox; BioBasic) for 18-24 h, or other concentrations as indicated and stable expression was confirmed with western blotting, as previously described (Zak and Antonescu, 2023).

### siRNA Transfection

siRNA transfection as performed as previously described (Orofiamma et al., 2025). Wild-type ARPE-19, pSBtet-BP-mSG-Gapex-5-WT, and pSBtet-BP-mSG-Gapex5-S902A cells lines were seeded on coverslips at 30% confluency. The next day, cells were transfected with either non-targeting siRNA, Gapex-5, or Hook3 siRNA sequences (found in **Table S1**) with Lipofectamine RNAiMAX (ThermoFisher). Each siRNA was precomplexed with the transfection reagent in Opti-MEM, incubated for 15 min at room temperature, and added dropwise to adhered cells to a final siRNA concentration of 220 pmol/L. Cells were incubated with the siRNA complexes for 4 h, followed by a PBS wash and replacement with full serum (10% FBS) DMEM/F12 for 24 hours.

### Fluorescence Staining

ARPE-19 cells were seeded onto glass coverslips in 6-well plates and cultured in DMEM/F12 supplemented with 10% FBS and transfected as indicated. For experiments involving rescue of endogenous Gapex-5 knockdown (**Figure 2D-E**), cells stably carrying a transgene for expression of either mSG-Gapex5 WT or S902A were transfected with Gapex-5-targeting siRNA (as described above), and the expression of the rescue construct was induced with 0.5 μM doxycycline for the last 24 h prior to labeling. Prior to fixation, cells were incubated in serum-free medium (DMEM/F12 + 0% FBS) for 1 h and subjected to treatments as indicated. For experiments requiring mitochondrial labeling, 200 nM MitoTracker Deep Red FM (ThermoFisher) was added during the final 30 minutes of incubation in serum-free media. In these cases, 4% paraformaldehyde (PFA) in DPBS was pre-warmed to 37C, and fixation was performed at room temperature for 20 min, protected from light. After fixation, cells were transferred to ice and washed three times with cold DPBS, followed by quenching with 100 mM glycine for 10 minutes and permeabilization in 0.1% Triton X-100 with 100 mM glycine for 10 minutes. For all other immunofluorescence experiments not involving MitoTracker, cells were fixed with ice-cold 4% PFA in DPBS for 20 min on ice, and all subsequent steps, including washes, quenching, and permeabilization, were likewise performed on ice. Following permeabilization, samples were blocked in 3% BSA in DPBS for 15 min and incubated with primary rabbit monoclonal anti-EEA1 or other primary antibody for 1 h at room temperature. After washing, cells were incubated with species-appropriate secondary antibodies conjugated to appropriate fluorophores. Nuclei were counterstained with DAPI (1 μg/mL) for 5 min before mounting in DAKO fluorescence mounting medium. Imaging was performed using spinning disc confocal microscope equipped with a 63× oil immersion objective (NA 1.65). Identical exposure settings were applied across conditions, and image analysis was conducted using FIJI (ImageJ, National Institutes of Health (Schneider et al., 2012)).

### Mitochondrial Iron Quantification and Live-Cell Imaging

ARPE-19 cells were seeded at 15% confluency on 18 mm glass coverslips in 12-well plates and cultured overnight in DMEM/F12 supplemented with 10% FBS at 37°C and 5% CO₂. The following day, cells were switched to serum-free DMEM/F12 medium and incubated for 4 h in media containing 6.25 μM human holo-Tfn (Sigma Aldrich), either in the presence of 100 μM A-769662 or vehicle control (0.1% v/v DMSO). Where indicated, deferoxamine (DFO, 10 mM) was added during the final hour as a negative control for mitochondrial iron detection. Additional experimental conditions included the use of a VPS34 inhibitor or transient transfection of ARPE-19 cells with siRNA or expression constructs.

To visualize mitochondrial iron (**Figure 5A-B**, **Figure 6**), cells were incubated with 5 μM Mito-FerroGreen (MFG, Dojindo) during the final 30 min of treatment, protected from light. In parallel experiments, MitoTracker Deep Red FM (200 nM, ThermoFisher) was used instead of MFG to label total mitochondrial mass (**Figure 5C-D**). Following incubation, coverslips were transferred to a live-cell imaging chamber and immediately imaged using a spinning disc confocal microscope equipped with a 63× oil immersion objective (NA 1.65). For each condition, 30 fields of view were acquired using matched exposure and laser settings. Images were analyzed using FIJI (ImageJ, National Institutes of Health (Schneider et al., 2012)), and fluorescence intensity was quantified from raw, unprocessed images.

### Tfn Internalization Assay

The internalization of fluorescent Tfn to early endosomes was performed as previously described (Bone et al., 2017). Following treatment with 100 μM A-7969662 for 15 min as indicated, cells were incubated with 20 μg/ml of A647-Tfn for indicated times at 37C and then immediately placed on ice and washed three times in ice-cold PBS2+ (PBS supplemented with 1mM Ca^2+^ and 1mM Mg^2+^) to remove unbound ligand, fixed in 4% paraformaldehyde, permeabilized in 0.1% Triton X-100, stained with anti-EEA1 and appropriate secondary antibodies, and then mounted in fluorescence mounting medium (Dako). Colocalization of A647-Tfn with EEA1 was performed in ImageJ using the Just Another Colocalization Plug-in (JaCoP) (Bolte and Cordelières, 2006) Pearson’s r-value.

### Tfn Recycling Assay

The internalization of fluorescent Tfn to early endosomes was performed based on a similar assay for measurement of recycling of the glucose transporter GLUT4 (Antonescu et al., 2008b). Cells were incubated with 20 μg/ml of A647-Tfn for 30 min at 37C in the presence or absence of 100 μM A-7969662 for the last 15 min as indicated. Cells were then washed in 37C media to remove A647-Tfn from the extracellular milieu, and then incubated in media lacking A647-Tfn for the indicated times. Cells were then immediately placed on ice and washed three times in ice-cold PBS to remove unbound ligand, fixed in 4% paraformaldehyde, and then mounted in fluorescence mounting medium (Dako). A647-Tfn intensity was measured by manual identification of cell outlines and measurement of mean fluorescence intensity using ImageJ. The loss of A647-Tfn signal from cells represents the fraction of Tfn/TfR that that undergone recycling from intracellular compartments to the plasma membrane.

### Whole-cell lysates and western blotting

Cell lysis and western blotting were performed as previously described (Cabral-Dias et al., 2022; Sugiyama et al., 2023; Rahmani et al., 2023; Orofiamma et al., 2025). After transfection or indicated treatments, cells were placed on ice and washed with PBS. Whole cell lysates were prepared in 2X LSB with a protease and phosphatase inhibitor cocktail (1LmM sodium orthovanadate, 10LnM okadaic acid, and 20LnM Protease Inhibitor Cocktail; all from BioShop). Lysates were supplemented with 10% β-mercaptoethanol and 5% bromophenol blue, heated at 65C for 15 min, and passed through a 27.5-gauge syringe. Proteins were resolved by glycine-Tris SDS–PAGE followed by transfer onto a 0.2 μm polyvinylidene fluoride membrane (Immobilon, Millipore). Membranes were blocked with 3% bovine serum albumin (BSA; BioShop) in TBST (Tris-buffered saline supplemented with Tween 20) and incubated with specific primary antibodies (**Table S1**) diluted in TBST supplemented with 1% BSA at 4C overnight. After TBST washes, membranes were incubated with the appropriate HRP-conjugated secondary antibodies (Cell Signaling Technology) (**Table S1**) at room temperature for 1 h. Following several TBST washes, bands were visualized using Luminata Crescendo HRP substrate (Millipore Sigma) on the Bio-Rad ChemiDoc Touch Imaging System.

### TMRE florescence measurement

ARPE19 cells were cultured in DMEM/F-12 supplemented with 10% FBS and 1% penicillin– streptomycin at 37 °C and 5% CO₂, seeded at 1.0 × 10L cells per well in 6-well plates, and treated the following day with 0.1% DMSO, 100 µM A-769662, or 10 µM FCCP for 2, 4, or 6 h. During the final 30 minutes of treatment, mitochondria were labeled with 0.1 µM TMRE. After 20 min of TMRE incubation, cells were washed three times with PBS, replenished with phenol-free DMEM, and imaged live on a Cytation 1 imager using consistent exposure and laser settings across conditions. Mitochondrial membrane potential (ΔΨm) was quantified as TMRE Integrated Density per cell on FIJI.

### Mitochondrial morphology analysis

Two-dimensional mitochondrial morphology analysis was performed to assess structural alterations following each treatment. For each condition, 30 individual cells were manually cropped from fluorescence images, and mitochondrial networks were analyzed using the Mitochondria Analyzer plugin in ImageJ/FIJI (Chaudhry et al., 2019), following the developer’s guidelines. Parameters such as mitochondrial total area, mean area, and branch length were extracted automatically. Mean values for all morphological parameters were calculated and compared across experimental conditions.

### Structural Illumination Microscopy (SIM)

ARPE19 cells were seeded on gridded coverslips at 1.0 × 10L cells per well in 6-well plates. On the day of the experiment (24h after seeding), cells were serum-starved for 1 hour, followed by incubation with Alexa Fluor 488–conjugated transferrin for 1 hour to label early endosomes. Mitochondria were subsequently stained with MitoTracker Deep Red (100 nM) for 20 min at 37C. Cells were then treated with either 0.01% DMSO or 100 µM A-769662 for 15 min and immediately fixed with 4% paraformaldehyde and 0.1% glutaraldehyde in PBS for 15 min at room temperature. Cells were washed three times with PBS and imaged on a Zeiss Elyra PS.1 Structured Illumination Microscope (SIM) with a Plan Apochromat 100× 1.46 NA objective lens. Five rotations of the grid pattern with three phases were collected for each z plane. Images were reconstructed with the SIM automatic setting module in Zeiss ZEN Black 2012 SP5 software.

### Statistical analysis

Statistical analyses were performed using GraphPad Prism 10 for MacOS, each statistical test is described in the figure captions.

