## Supplemental Material for "AMPK Repositions Early Endosomes via Gapex-5 to Promote Delivery of Iron to Mitochondria"

This supplemental material contains the following:

**Table S1**: Details of reagents used in this study
**Figure S1**: mTORC1 inhibition does not trigger EEA1 early endosome repositioning
**Figure S2**: Silencing and rescue of Gapex-5 expression
**Figure S3**: Hook3 silencing
**Figure S4**: Treatment with A-769662 does not alter overlap of early endosomes and lysosomes
**Figure S5**: Treatment with A-769662 increases TMRE fluorescence and alters mitochondrial morphology

| Target | Sense | Anti-sense |
|---|---|---|
| GAPVD1 (#1) | N/A (Horizon Discovery, J-026206-12) | N/A (Horizon Discovery, J-026206-12) |
| GAPVD1 (#2) | CCA ACA GGC GAG UGG CGA A(UU) | AAU UCG CCA CUC GCC UGU UGG |
| HOOK3 | GGA GAU AAU UGG AGG CUA A (UU) | UUA GCC UCC AAU UAU CUC C(UU) |
| Control siRNA (Con7) | CGU ACU GCU UGC GAU ACG GUU | CCG UAU CGC AAG CAG UAC GUU |

| | Company | Product Code |
|---|---|---|
| **Immunofluorescence Antibodies** | | |
| EEA1 (E9Q6G) | Cell Signaling Technology | 48453S |
| Lamp1 (D2D11) | Cell Signaling Technology | 9091S |
| Cy™3 AffiniPure® Donkey Anti-Mouse IgG (H+L) (min X Bov, Ck, Gt, GP, Sy Hms, Hrs, Hu, Rb, Rat, Shp Sr Prot) | Jackson ImmunoResearch | |
| **Western Blotting Antibodies** | | |
| GAPVD1 Polyclonal Antibody | Proteintech | 16940-1-AP |
| HOOK3 Polyclonal Antibody | Proteintech | 15457-1-AP |
| Pan-actin Rabbit mAb | Cell Signalling | D18c11 |

**Table S1.** Details of reagents used in this study, including sequences for siRNA gene silencing, and antibodies used for immunofluorescence and Western blotting.

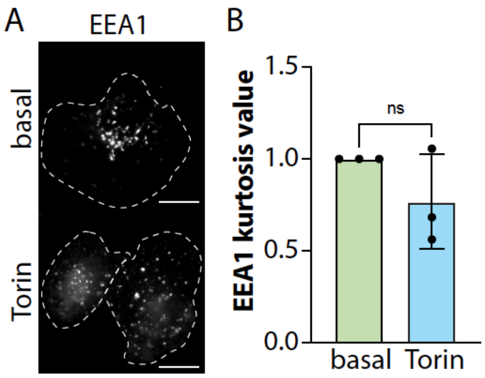


**Figure S1. mTORC1 inhibition does not trigger EEA1 early endosome repositioning.** ARPE-19 cells were treated with Torin or vehicle control. Shown are representative micrographs obtained by spinning disc confocal microscopy, scale 20 μm (A) and quantification of kurtosis value, shown as mean ± SD from 3 independent experiments (normalized to basal) (B), with each experiment involving measurement of at least 30 cells per condition. Statistical comparisons were performed by Mann-Whitney test.

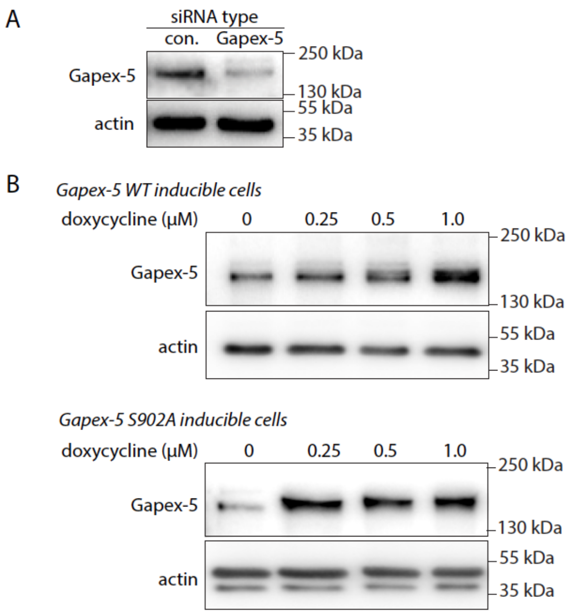


**Figure S2. Silencing and rescue of Gapex-5 expression**. (***A***) ARPE-19 cells were subjected to siRNA silencing using sequences targeting Gapex-5 or non-targeting control (con.), as in **Figure 2A-B**. (***B***) Stable cells engineered using the Sleeping Beauty transposon system that carry transgenes for inducible expression of Gapex-5 WT or S902A were treated with doxycycline as indicated for 24h (as in **Figure 2C-D**). For all, whole cell lysates were resolved by immunoblotting, using antibodies specific for Gapex-5 or actin. In (B), Gapex-5 S902A appears to induce higher levels of expression at given doxycycline concentrations than WT Gapex-5. For all knockdown-rescue experiments, 0.5 μM doxycycline was used.

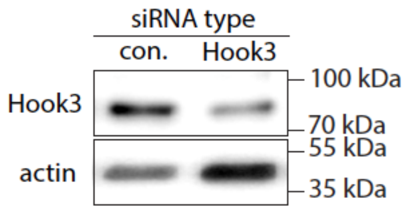


**Figure S3. Hook3 silencing**. ARPE-19 cells were subjected to siRNA silencing using sequences targeting Hook3 or non-targeting control (con.), as in **Figure 3A-B**. Whole cell lysates were resolved by immunoblotting, using antibodies specific for Hook3 or actin.

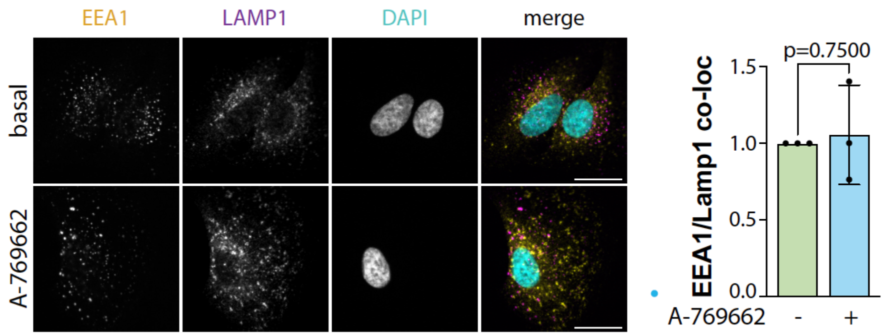


**Figure S4. Treatment with A-769662 does not alter overlap of early endosomes and lysosomes**. ARPE-19 cells were treated with 100 μm A769662 or vehicle (0.1% v/v DMSO) control for 15 min. Shown (left panels) are representative images obtained by spinning disc confocal microscopy, scale 20 μm, and (right panels) quantification of co-localization between EEA1 and LAMP1 by Manders' coefficient, shown as mean ± SD from 3 independent experiments, with each experiment involving measurement of at least 30 cells per condition (normalized to untreated condition). Statistical comparisons were performed by Mann-Whitney test.

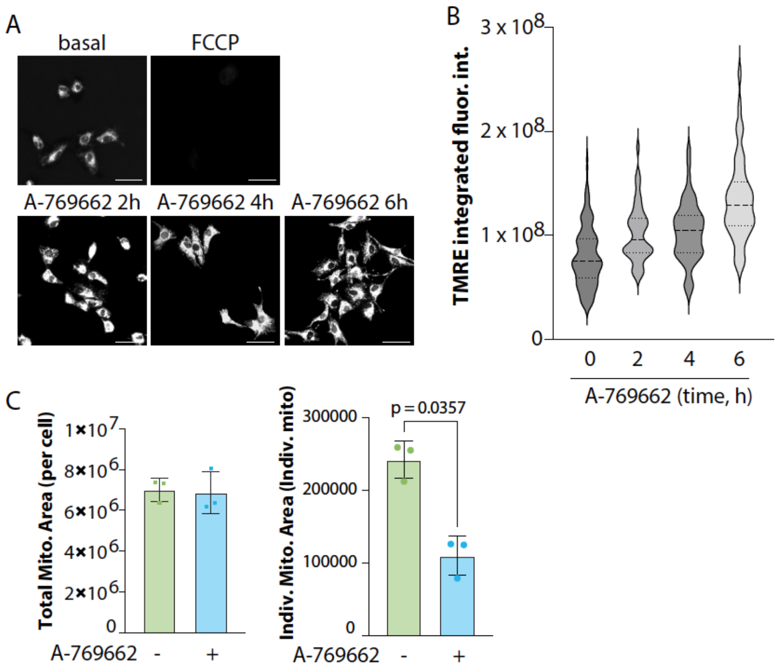


**Figure S5. Treatment with A-769662 increases TMRE fluorescence and alters mitochondrial morphology**. (***A-B***) ARPE-19 cells were treated with 100 μM A-769662 and then subjected to labeling with TMRE, followed by detection of TMRE fluorescence using the Cytation imaging platform. Shown are representative fluorescence images of the TMRE fluorescence, scale 50 μm (A) and quantification of mean TMRE fluorescence intensity in individual cells (B). (***C***) Fluorescence images corresponding to cells labelled with MitoTracker (**Figure 5C-D**) were subject to mitochondrial morphology analysis. Shown for the total mitochondrial area per cell and the mean mitochondrial area (area of individual mitochondria) as mean ± SD from 3 independent experiments, with each experiment involving measurement of at least 30 cells per condition. Statistical comparisons were performed by paired t-test.
